# Linked emergence of racial disparities in mental health and epigenetic biological aging across childhood and adolescence

**DOI:** 10.1101/2024.03.26.586786

**Authors:** Muna Aikins, Yayouk Willems, Colter Mitchell, Bridget Goosby, Laurel Raffington

## Abstract

Marginalization due to structural racism may confer an increased risk for aging-related diseases – in part – via effects on people’s mental health. Here we leverage a prospective birth cohort study to examine whether the emergence of racial disparities in mental health and DNA-methylation measures of biological aging (*i*.*e*., DunedinPACE, GrimAge Acceleration, PhenoAge Acceleration) are linked across childhood and adolescence. We further consider to what extent racial disparities are statistically accounted for by perinatal and postnatal factors in preregistered analyses of N=4,898 participants from the Future of Families & Child Wellbeing Study, of which N=2,039 had repeated saliva DNA methylation at ages 9 and 15 years. We find that racially marginalized children had higher levels of externalizing and internalizing behaviors and diverging longitudinal internalizing slopes. Black compared to White identifying children, children living in more racially segregated neighborhoods, and racially marginalized children more affected by colorism tended to have higher age-9 levels of biological aging and more biological age acceleration over adolescence. Notably, longitudinal increases in internalizing and externalizing behavior were correlated with longitudinal increases in biological aging. While racial and ethnic disparities in mental health were largely statistically accounted for by socioeconomic variables, racial differences in biological aging were often still visible beyond covariate controls. Our findings indicate that racial disparities in mental health and biological aging are linked and emerge early in life. Programs promoting racial health equity must address the psychological and physical impacts of structural racism in children. Comprehensive measures of racism are lacking in current population cohorts.

## Introduction

A large body of evidence has recorded striking racial disparities in physical and mental health (Arrondo et al., 2022; Bailey et al., 2021; Creanga, 2018; Krieger, 2021; Pachter & Coll, 2009). Therefore, examining how different manifestations of racism, including effects of institutionalized systems and interpersonal social dynamics in which individuals are “racialized”, affects health across the lifespan is a priority to improving population health (Acker et al., 2023; Iruka et al., 2022; Williams et al., 2019). For instance, heightened daily life stress and vigilance stemming from ongoing racialization may amplify the risk of lower mental health and contribute to higher levels of chronic inflammation and accelerated multisystem biological aging (Castro-Ramirez et al., 2021; Cheadle et al., 2020; Deckard et al., 2023; Goosby et al., 2018; Jochman et al., 2019; Poganik et al., 2023). Biological aging can be defined as the progressive loss of system integrity that occurs with advancing chronological age, including changes in DNA-methylation (DNAm; Horvath & Raj, 2018; Kirkwood, 2005). DNAm measures of biological aging can be applied early in the life course to study the etiology of social health disparities, decades before differences in disease and mortality are measurable (Raffington & Belsky, 2022).

Racial marginalization and low socioeconomic status has been associated with more advanced and faster biological aging as measured in DNAm in both adults and children, and, in adults, these differences in biological aging partially account for health disparities between and within racial groups (Del Toro et al., 2022; Oblak et al., 2021; Raffington et al., 2021; Sugden et al., 2023). A few studies have found DNAm measures of biological aging to be associated with mental health (Cecil et al., 2023; Oblak et al., 2021; Raffington, Tanksley, et al., 2023). Yet, previous research has largely been cross-sectional in design and, thus, does not address the dynamic interplay between racialization, mental health, and biological aging.

Here we leverage a prospective birth cohort study to examine whether the emergence of racial disparities in mental health is linked to racial disparities in DNA-methylation measures of biological aging across childhood and adolescence. We further consider to what extent racial disparities are statistically accounted for by perinatal (*e*.*g*., birthweight) and postnatal (*e*.*g*., socioeconomic status, body mass index) factors in preregistered analyses of N=4,898 participants from the Future of Families & Child Wellbeing Study, which intentionally oversampled financially under resourced families. Among the N=2,039 participants who had repeated saliva DNAm at ages 9 and 15 years, most participants racially positioned themselves as African-American/Black (n=901, 47%), followed by Hispanic/Latinx (n=511, 26%), White (n=366, 19%), Multiracial (n=99, 5%), and “Other” (n=52, 3%).

## Results

Our preregistered analyses (https://osf.io/xbgzu) and results are categorized into three objectives:

1. We examined racial and ethnic disparities in parent-reported internalizing and externalizing behaviors in longitudinal growth curve models from early childhood through adolescence and in cross-sectional analyses of self-reported anxiety and depressive symptoms at age 15. Racial and ethnic disparities were conceptualized as outcomes of structural racism, more specifically, as “racialization”, which emphasizes the social processes and institutionalized systems in which individuals are positioned (Bonilla-Silva, 1997; Powell, 2008, 2013). We quantified measures of race as a self-identified social position, the Thiel Index of racial neighborhood segregation, and – amongst marginalized youth only – skin tone as a proxy of colorism that serves to maintain the racial hierarchy (Bailey et al., 2017, 2021; Dixon & Telles, 2017; Hunter, 2007; Monk, 2021; Roberto, 2016)
2. Next, we tested for racial and ethnic disparities in saliva DNAm quantifications of biological aging (DunedinPACE, GrimAge Acceleration, PhenoAge Acceleration), combining repeated DNAm measures from ages 9 and 15 in univariate latent change score models. While these DNAm measures of biological aging were developed in blood (Belsky et al., 2022; Levine et al., 2018; Lu et al., 2019), our previous findings suggest good cross-tissue correspondence to saliva DNAm residualized for cell composition, presumably because of the high immune cell signal in both blood and saliva (Middleton et al., 2022; Raffington, Schneper, et al., 2023a; Raffington, Tanksley, et al., 2023).
3. Lastly, we assessed if changes in internalizing and externalizing behaviors from 9-to-15-years were correlated with changes in biological aging from 9-to-15-years by applying bivariate latent change score models (Kievit et al., 2018; McArdle, 2009). We report nominal *p*-values with an alpha <.05 threshold and note if results remain significant after Benjamini-Hochberg False-Discovery-Rate correction (FDR, Benjamini & Hochberg, 1995). See Table 2 for a description of measures, **Supplemental Material Table 1** for a list of preregistered analyses and analytic deviations, and **Supplemental Table 1** for descriptive statistics and correlations between measures.

**Table 1.** Longitudinal changes in externalizing and internalizing behaviors with longitudinal changes in biological aging from 9-to-15-years.

| BioAge | Parameters | Externalizing |  |  |  |  | Internalizing |  |  |  |  |
| --- | --- | --- | --- | --- | --- | --- | --- | --- | --- | --- | --- |
|  |  | *r* | SE | *p* | 95% CI |  | *r* | SE | *p* | 95% CI |  |
| DunedinPACE | MH_I & BioAge_I | .02 | .02 | .48 | -.03 | .06 | .04 | .02 | .12 | -.01 | .09 |
|  | MH_I & BioAge_LC | **.07** | **.02** | **.00** | **.03** | **.12** | .03 | .02 | .20 | -.02 | .08 |
|  | BioAge_I & MH_LC | -.01 | .02 | .66 | -.06 | .04 | -.03 | .02 | .22 | -.07 | .02 |
|  | MH_LC & BioAge_LC | **.06** | **.02** | **.01** | **.03** | **.13** | **.06** | **.02** | **.01** | **.01** | **.10** |
| GrimAge Acceleration | MH_I & BioAge_I | **.05** | **.02** | **.04** | **.01** | **.09** | **.06** | **.02** | **.01** | **.02** | **.11** |
|  | MH_I & BioAge_LC | **.05** | **.02** | **.04** | **.01** | **.09** | -.01 | .03 | .59 | -.06 | .04 |
|  | BioAge_I & MH_LC | .02 | .02 | .45 | -.03 | .06 | **-.06** | **.02** | **.01** | **-.11** | **-.02** |
|  | MH_LC & BioAge_LC | **.06** | **.02** | **.01** | **.02** | **.11** | .04 | .02 | .10 | -.01 | .08 |
| PhenoAge Acceleration | MH_I & BioAge_I | **.06** | **.02** | **.01** | **.01** | **.11** | .03 | .02 | .20 | -.02 | .08 |
|  | MH_I & BioAge_LC | .02 | .02 | .41 | -.03 | .07 | -.06 | .02 | .02 | -.11 | -.01 |
|  | BioAge_I & MH_LC | .02 | .02 | .47 | -.03 | .06 | -.06 | .02 | .01 | -.10 | -.02 |
|  | MH_LC & BioAge_LC | **.05** | **.02** | **.02** | **.01** | **.10** | **.06** | **.02** | **.01** | **.01** | **.10** |

Note: Parameters correspond to path labels in Figure 3, *MH\_I & BioAge\_I* = correlation between mental health intercept and biological age intercept, *MH\_I & BioAge\_LC* = correlation between Intercept mental health and longitudinal change in *BioAge*, *BioAge\_I & MH\_LC* = correlation between Intercept DNAm and longitudinal change in mental health, *MH\_LC & BioAge\_LC* = correlation between longitudinal change in mental health and longitudinal change in *BioAge*. Significant associations at $p < .05$ are marked in bold. These associations hold after FDR correction.

**Table 2.**
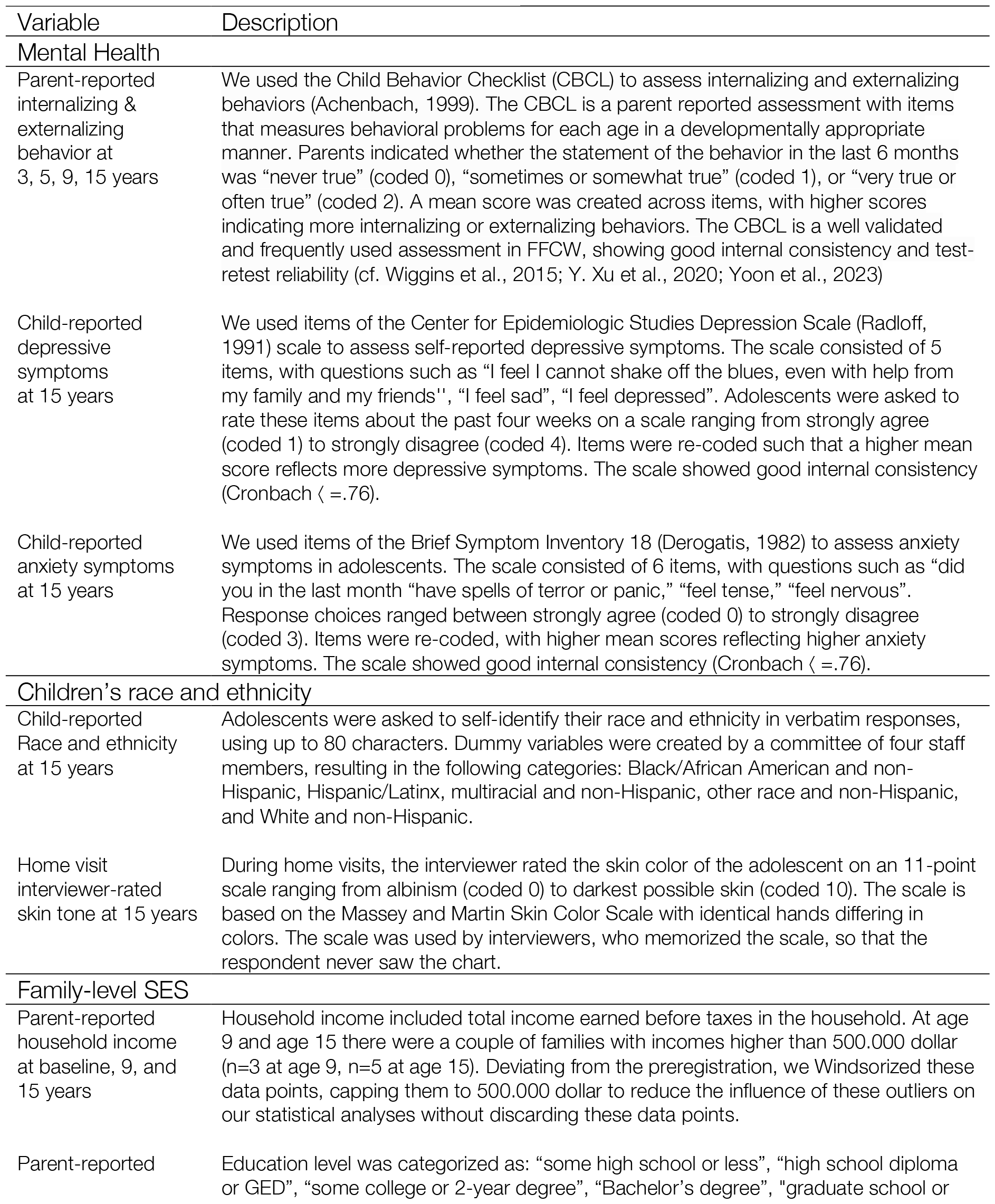
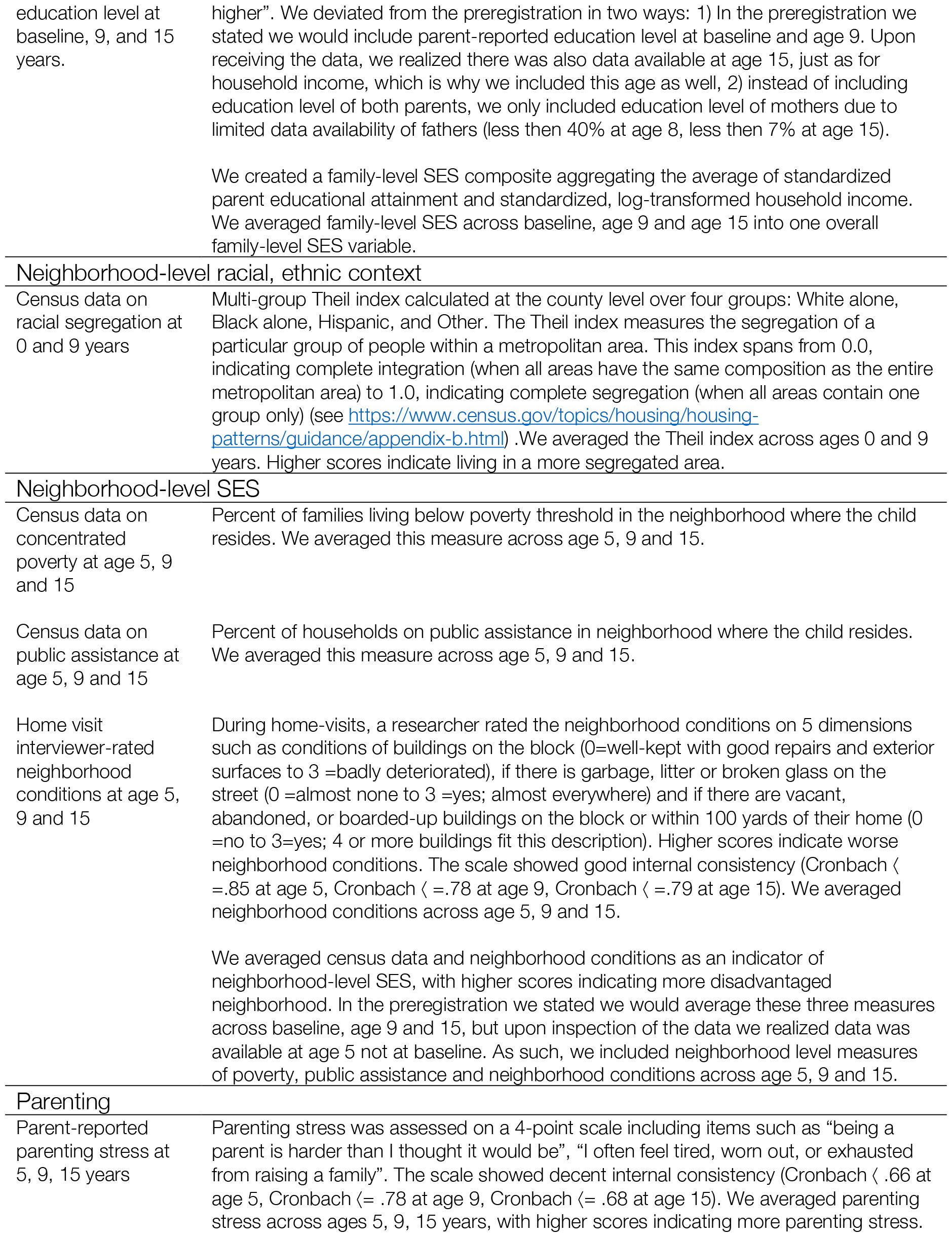
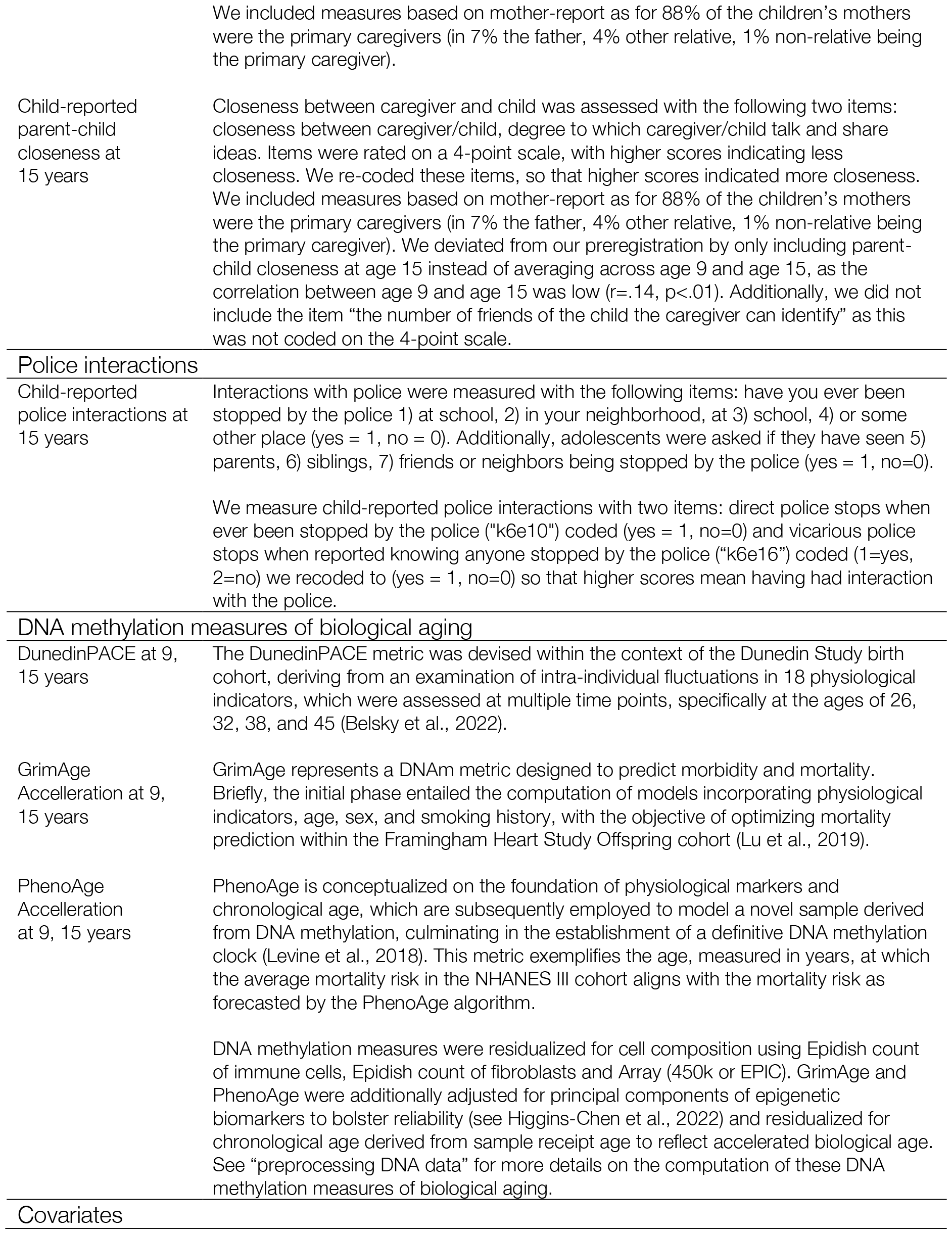
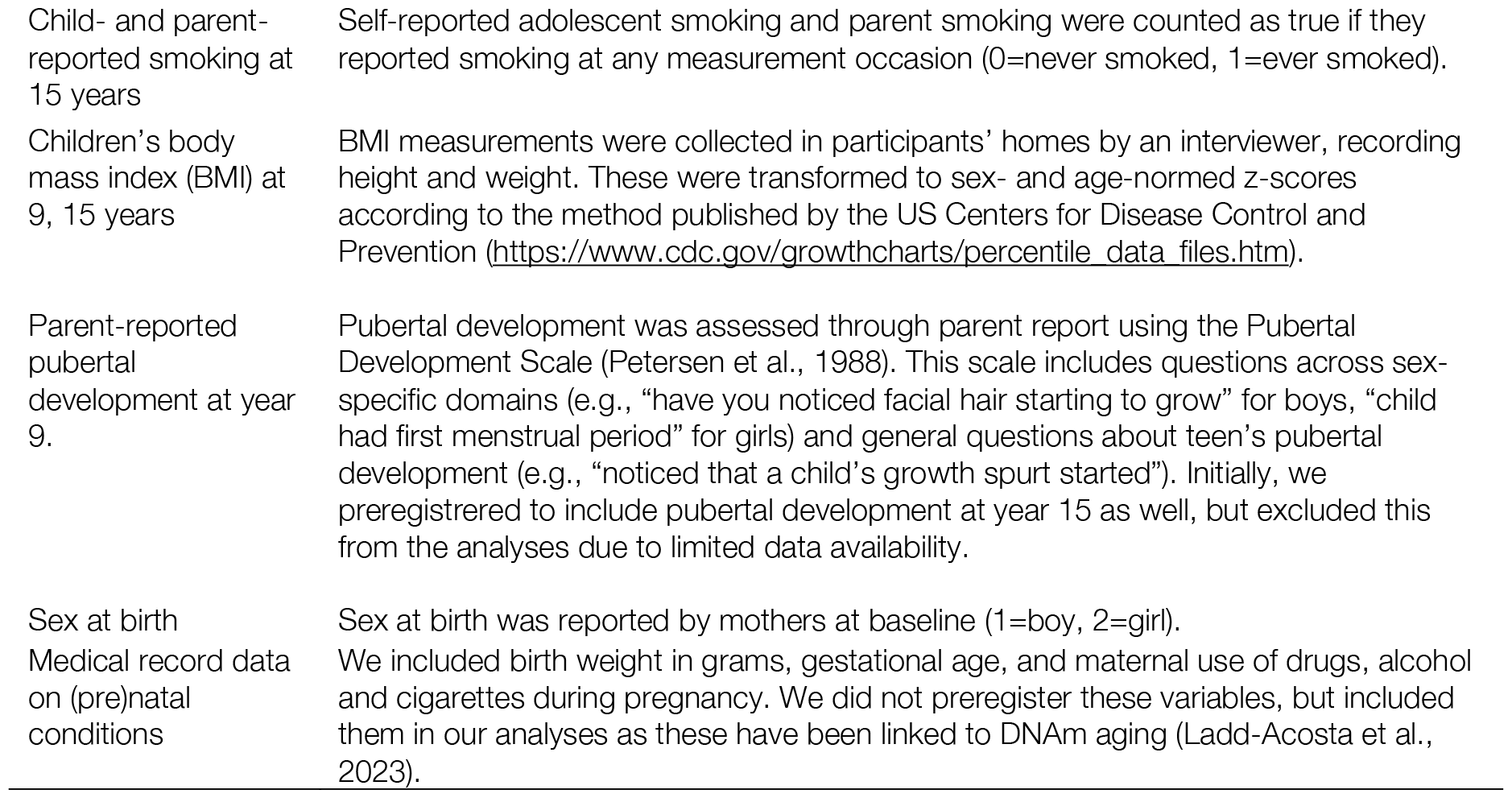
Description of measures.

| Variable | Description |
| --- | --- |
| **Mental Health** |  |
| Parent-reported internalizing & externalizing behavior at 3, 5, 9, 15 years | We used the Child Behavior Checklist (CBCL) to assess internalizing and externalizing behaviors (Achenbach, 1999). The CBCL is a parent reported assessment with items that measures behavioral problems for each age in a developmentally appropriate manner. Parents indicated whether the statement of the behavior in the last 6 months was “never true” (coded 0), “sometimes or somewhat true” (coded 1), or “very true or often true” (coded 2). A mean score was created across items, with higher scores indicating more internalizing or externalizing behaviors. The CBCL is a well validated and frequently used assessment in FFCW, showing good internal consistency and test-retest reliability (cf. Wiggins et al., 2015; Y. Xu et al., 2020; Yoon et al., 2023) |
| Child-reported depressive symptoms at 15 years | We used items of the Center for Epidemiologic Studies Depression Scale (Radloff, 1991) scale to assess self-reported depressive symptoms. The scale consisted of 5 items, with questions such as “I feel I cannot shake off the blues, even with help from my family and my friends”, “I feel sad”, “I feel depressed”. Adolescents were asked to rate these items about the past four weeks on a scale ranging from strongly agree (coded 1) to strongly disagree (coded 4). Items were re-coded such that a higher mean score reflects more depressive symptoms. The scale showed good internal consistency (Cronbach $\alpha = .76$). |
| Child-reported anxiety symptoms at 15 years | We used items of the Brief Symptom Inventory 18 (Derogatis, 1982) to assess anxiety symptoms in adolescents. The scale consisted of 6 items, with questions such as “did you in the last month “have spells of terror or panic,” “feel tense,” “feel nervous”. Response choices ranged between strongly agree (coded 0) to strongly disagree (coded 3). Items were re-coded, with higher mean scores reflecting higher anxiety symptoms. The scale showed good internal consistency (Cronbach $\alpha = .76$). |
| **Children’s race and ethnicity** |  |
| Child-reported Race and ethnicity at 15 years | Adolescents were asked to self-identify their race and ethnicity in verbatim responses, using up to 80 characters. Dummy variables were created by a committee of four staff members, resulting in the following categories: Black/African American and non-Hispanic, Hispanic/Latinx, multiracial and non-Hispanic, other race and non-Hispanic, and White and non-Hispanic. |
| Home visit interviewer-rated skin tone at 15 years | During home visits, the interviewer rated the skin color of the adolescent on an 11-point scale ranging from albinism (coded 0) to darkest possible skin (coded 10). The scale is based on the Massey and Martin Skin Color Scale with identical hands differing in colors. The scale was used by interviewers, who memorized the scale, so that the respondent never saw the chart. |
| **Family-level SES** |  |
| Parent-reported household income at baseline, 9, and 15 years | Household income included total income earned before taxes in the household. At age 9 and age 15 there were a couple of families with incomes higher than 500,000 dollar ($n=3$ at age 9, $n=5$ at age 15). Deviating from the preregistration, we Winsorized these data points, capping them to 500,000 dollar to reduce the influence of these outliers on our statistical analyses without discarding these data points. |
| Parent-reported | Education level was categorized as: “some high school or less”, “high school diploma or GED”, “some college or 2-year degree”, “Bachelor’s degree”, “graduate school or |
| education level at baseline, 9, and 15 years. | higher". We deviated from the preregistration in two ways: 1) In the preregistration we stated we would include parent-reported education level at baseline and age 9. Upon receiving the data, we realized there was also data available at age 15, just as for household income, which is why we included this age as well, 2) instead of including education level of both parents, we only included education level of mothers due to limited data availability of fathers (less than 40% at age 8, less than 7% at age 15).<br>We created a family-level SES composite aggregating the average of standardized parent educational attainment and standardized, log-transformed household income. We averaged family-level SES across baseline, age 9 and age 15 into one overall family-level SES variable. |
| **Neighborhood-level racial, ethnic context** |  |
| Census data on racial segregation at 0 and 9 years | Multi-group Theil index calculated at the county level over four groups: White alone, Black alone, Hispanic, and Other. The Theil index measures the segregation of a particular group of people within a metropolitan area. This index spans from 0.0, indicating complete integration (when all areas have the same composition as the entire metropolitan area) to 1.0, indicating complete segregation (when all areas contain one group only) (see https://www.census.gov/topics/housing/housing-patterns/guidance/appendix-b.html). We averaged the Theil index across ages 0 and 9 years. Higher scores indicate living in a more segregated area. |
| **Neighborhood-level SES** |  |
| Census data on concentrated poverty at age 5, 9 and 15 | Percent of families living below poverty threshold in the neighborhood where the child resides. We averaged this measure across age 5, 9 and 15. |
| Census data on public assistance at age 5, 9 and 15 | Percent of households on public assistance in neighborhood where the child resides. We averaged this measure across age 5, 9 and 15. |
| Home visit interviewer-rated neighborhood conditions at age 5, 9 and 15 | During home-visits, a researcher rated the neighborhood conditions on 5 dimensions such as conditions of buildings on the block (0=well-kept with good repairs and exterior surfaces to 3=badly deteriorated), if there is garbage, litter or broken glass on the street (0 =almost none to 3 =yes; almost everywhere) and if there are vacant, abandoned, or boarded-up buildings on the block or within 100 yards of their home (0 =no to 3=yes; 4 or more buildings fit this description). Higher scores indicate worse neighborhood conditions. The scale showed good internal consistency (Cronbach $\alpha$ =.85 at age 5, Cronbach $\alpha$ =.78 at age 9, Cronbach $\alpha$ =.79 at age 15). We averaged neighborhood conditions across age 5, 9 and 15.<br>We averaged census data and neighborhood conditions as an indicator of neighborhood-level SES, with higher scores indicating more disadvantaged neighborhood. In the preregistration we stated we would average these three measures across baseline, age 9 and 15, but upon inspection of the data we realized data was available at age 5 not at baseline. As such, we included neighborhood level measures of poverty, public assistance and neighborhood conditions across age 5, 9 and 15. |
| **Parenting** |  |
| Parent-reported parenting stress at 5, 9, 15 years | Parenting stress was assessed on a 4-point scale including items such as "being a parent is harder than I thought it would be", "I often feel tired, worn out, or exhausted from raising a family". The scale showed decent internal consistency (Cronbach $\alpha$ =.66 at age 5, Cronbach $\alpha$ =.78 at age 9, Cronbach $\alpha$ =.68 at age 15). We averaged parenting stress across ages 5, 9, 15 years, with higher scores indicating more parenting stress. |
|  | We included measures based on mother-report as for 88% of the children's mothers were the primary caregivers (in 7% the father, 4% other relative, 1% non-relative being the primary caregiver). |
| Child-reported parent-child closeness at 15 years | Closeness between caregiver and child was assessed with the following two items: closeness between caregiver/child, degree to which caregiver/child talk and share ideas. Items were rated on a 4-point scale, with higher scores indicating less closeness. We re-coded these items, so that higher scores indicated more closeness. We included measures based on mother-report as for 88% of the children's mothers were the primary caregivers (in 7% the father, 4% other relative, 1% non-relative being the primary caregiver). We deviated from our preregistration by only including parent-child closeness at age 15 instead of averaging across age 9 and age 15, as the correlation between age 9 and age 15 was low ($r=.14$, $p<.01$). Additionally, we did not include the item "the number of friends of the child the caregiver can identify" as this was not coded on the 4-point scale. |
| **Police interactions** |  |
| Child-reported police interactions at 15 years | Interactions with police were measured with the following items: have you ever been stopped by the police 1) at school, 2) in your neighborhood, at 3) school, 4) or some other place (yes = 1, no = 0). Additionally, adolescents were asked if they have seen 5) parents, 6) siblings, 7) friends or neighbors being stopped by the police (yes = 1, no=0).<br>We measure child-reported police interactions with two items: direct police stops when ever been stopped by the police ("k6e10") coded (yes = 1, no=0) and vicarious police stops when reported knowing anyone stopped by the police ("k6e16") coded (1=yes, 2=no) we recoded to (yes = 1, no=0) so that higher scores mean having had interaction with the police. |
| **DNA methylation measures of biological aging** |  |
| DunedinPACE at 9, 15 years | The DunedinPACE metric was devised within the context of the Dunedin Study birth cohort, deriving from an examination of intra-individual fluctuations in 18 physiological indicators, which were assessed at multiple time points, specifically at the ages of 26, 32, 38, and 45 (Belsky et al., 2022). |
| GrimAge Acceleration at 9, 15 years | GrimAge represents a DNAm metric designed to predict morbidity and mortality. Briefly, the initial phase entailed the computation of models incorporating physiological indicators, age, sex, and smoking history, with the objective of optimizing mortality prediction within the Framingham Heart Study Offspring cohort (Lu et al., 2019). |
| PhenoAge Acceleration at 9, 15 years | PhenoAge is conceptualized on the foundation of physiological markers and chronological age, which are subsequently employed to model a novel sample derived from DNA methylation, culminating in the establishment of a definitive DNA methylation clock (Levine et al., 2018). This metric exemplifies the age, measured in years, at which the average mortality risk in the NHANES III cohort aligns with the mortality risk as forecasted by the PhenoAge algorithm.<br>DNA methylation measures were residualized for cell composition using EpiDish count of immune cells, EpiDish count of fibroblasts and Array (450k or EPIC). GrimAge and PhenoAge were additionally adjusted for principal components of epigenetic biomarkers to bolster reliability (see Higgins-Chen et al., 2022) and residualized for chronological age derived from sample receipt age to reflect accelerated biological age. See "preprocessing DNA data" for more details on the computation of these DNA methylation measures of biological aging. |
| **Covariates** |  |
| Child- and parent-reported smoking at 15 years | Self-reported adolescent smoking and parent smoking were counted as true if they reported smoking at any measurement occasion (0=never smoked, 1=ever smoked). |
| Children's body mass index (BMI) at 9, 15 years | BMI measurements were collected in participants' homes by an interviewer, recording height and weight. These were transformed to sex- and age-normed z-scores according to the method published by the US Centers for Disease Control and Prevention (https://www.cdc.gov/growthcharts/percentile_data_files.htm). |
| Parent-reported pubertal development at year 9. | Pubertal development was assessed through parent report using the Pubertal Development Scale (Petersen et al., 1988). This scale includes questions across sex-specific domains (e.g., "have you noticed facial hair starting to grow" for boys, "child had first menstrual period" for girls) and general questions about teen's pubertal development (e.g., "noticed that a child's growth spurt started"). Initially, we preregistered to include pubertal development at year 15 as well, but excluded this from the analyses due to limited data availability. |
| Sex at birth | Sex at birth was reported by mothers at baseline (1=boy, 2=girl). |
| Medical record data on (pre)natal conditions | We included birth weight in grams, gestational age, and maternal use of drugs, alcohol and cigarettes during pregnancy. We did not preregister these variables, but included them in our analyses as these have been linked to DNAm aging (Ladd-Acosta et al., 2023). |

### (1) Racial and ethnic disparities in mental health from early childhood through adolescence

First, we examined associations of self-identified racial/ethnic positions, neighborhood segregation, and skin tone with externalizing behaviors in the full sample of N=4,898. We found that Black and Multiracial children compared to White identifying children had a higher externalizing intercept across early childhood through adolescence (Black: b= .09, 95% CI = .03 to .16, *p* <.01, Multiracial: b= .06, 95% CI=.01 to .0,11, *p*<.05, significant after FDR correction). Black compared to White children showed a steeper decline over childhood (b= -.11, CI = -.21 to -.01, *p*<.05), and a steeper increase across adolescence (b= .18, CI = .03 to 0.3, *p*<.05), but these differences did not survive FDR correction for multiple comparisons (**Figure 1 B; Supplemental Table 3**). A race by gender interaction suggested this decline over early childhood and increase across adolescence were particularly pronounced for Black boys, but this interaction did not remain after FDR correction (**Supplemental Table 3**).

**Figure 1.**
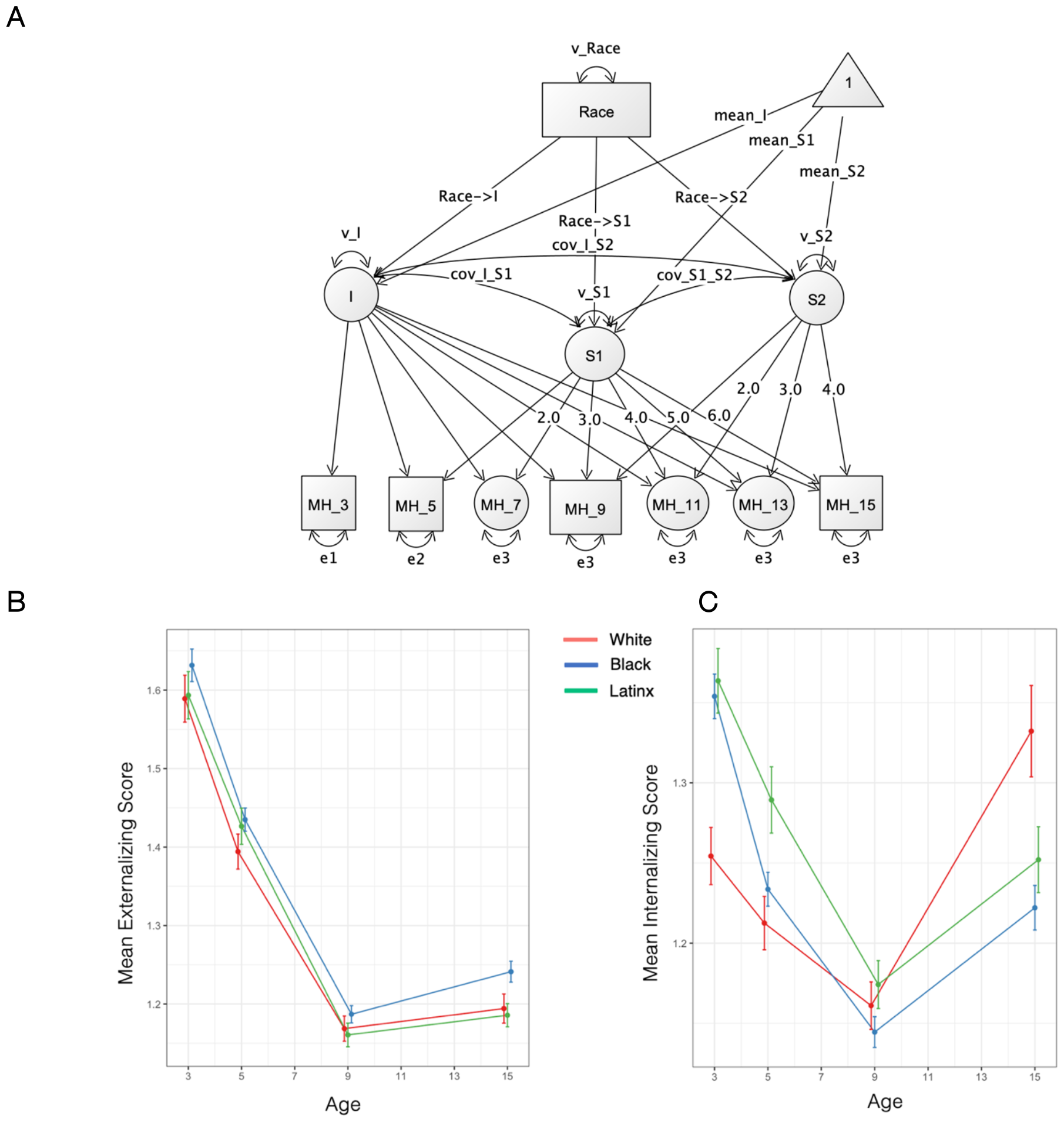
Racial and ethnic disparities in internalizing and externalizing behaviors from early childhood through adolescence. Panel. **A** depicts a latent growth curve model of mental health and race. Squares represent observed variables. Circles represent latent factors, including missing waves, intercepts, and slopes. Triangles denote constants, i.e. mean intercept and slopes. Single headed arrows denote regressions and double headed arrows denote covariances. **Panel B** and **C** depict mean scores in parent-reported externalizing and internalizing behaviors, respectively, for White, Black, and Latinx children. Data for the smaller subsamples of Multiracial and Other are not shown for visualization purposes but were included in the analyses.

Further, children living in more racially segregated neighborhoods, (who were more likely to identify as Black and Multiracial, **Supplemental Material Figure 1**), had a higher externalizing intercept (b=.09, CI = .04 to .13, *p*< .05, significant after FDR correction) and a steeper decline over early childhood (b= -.08, CI = -.15 to .01, *p*< .05, significant after FDR correction, **Supplemental Table 5**). Amongst Black, Latinx and Multiracial children, darker skin tone was associated with a higher externalizing intercept (b= .09, CI = .01 to .18, *p*< .05, significant after FDR correction, **Supplemental Table 6**).

Second, we examined main associations of racial/ethnic identities, neighborhood segregation, and skin tone with internalizing behaviors. We found that all groups of racially marginalized children showed a higher internalizing intercept across early childhood through adolescence compared to White children (Black b= .27, 95% CI = .20 to .34, *p* < .01; Latinx b = .28, CI = .21 to .35, *p* < .01; Other b = .06, CI = .01 to .12, *p*< .01; Multiracial b = .12, CI = .06 to .18, *p*< .01, significant after FDR correction, **Figure 1C, Supplemental Table 4**). They also showed a steeper decrease in internalizing behavior across early childhood compared to White children (Black b=-.34, 95% CI = -.48. to -.20, *p* < .01; Latinx b= -.21, CI = -.33 to -.09, *p* < .01; other race b= -.10, CI = -.20 to -.01, *p*< .05; Multiracial b=-.12, CI = -.22 to -.02, *p*< .05, all significant after FDR correction). A race by gender interaction showed that Black compared to White boys had a steeper increase in internalizing across adolescence (b= .23, 95%Cl= .08 to .38, *p*<.01, significant after FDR correction, **Figure 1C, Supplemental Table 4**).

Further, children living in more racially segregated neighborhoods had a higher internalizing intercept (b= .07, CI = .02 to .12, *p*< .01, significant after FDR correction), and a steeper decline in internalizing behavior across childhood (b=-.12, CI =-0.20 to -.04, *p*< .01, significant after FDR correction, **Supplemental Table 5**). Amongst Black, Latinx and Multiracial children, darker skin tone was not significantly associated with internalizing behaviors (**Supplemental Table 6**).

Next, we assessed to what extent these racial/ethnic disparities in child mental health were statistically accounted for by covariate adjustment for proximal contextual factors related to structural racialization (*e*.*g*., family socioeconomic status, neighborhood socioeconomic disadvantage, police interactions, parenting stress and closeness). Racial and ethnic disparities in externalizing and internalizing behaviors were largely statistically accounted for by covariate control for family socioeconomic status and neighborhood socioeconomic disadvantage, whereas covariate control for police interactions and parenting had little effect (**Supplemental Tables 3-6**). Importantly, all groups of racially marginalized children were far more likely to live in socioeconomically under resourced families and neighborhoods, whereas age-15 adolescent reports of police interactions and parenting stress showed racialized differences for Black compared to White children only (**Supplemental Table 2 and Supplemental Material Figure 1**).

We did not find evidence of racial/ethnic disparities in self-reported anxiety or depressive symptoms at age 15 (**Supplemental table 7**). Amongst marginalized youth only, darker skin tone was associated with more anxiety symptoms (b=-.07, 95% CI = -.14 to -.01, *p*<.05), but this result did not survive FDR correction (**Supplemental table 7**).

### (2) Racial and ethnic disparities in biological aging across adolescence

We tested for associations of racial/ethnic identities, neighborhood segregation, and skin tone with biological aging measured at age 9 and 15 years in N=2,039 children with DNAm (see **Figure 3** for a graphical model illustration). We found that Black and Latinx compared to White youth tended to have a higher intercept and higher longitudinal increase in the pace of aging (DunedinPACE) across adolescence (**Figure 2A & B, Supplemental Table 8**, significant after FDR correction). Black compared to White children also had a more advanced biological age intercept and higher longitudinal increase in biological age (GrimAge and PhenoAge Acceleration; **Supplemental Table 8, Figure 2**, significant after FDR correction). Further, children living in more racially segregated neighborhoods had a higher intercept in all measures of biological aging, and a higher longitudinal increase as indicated by DunedinPACE and PhenoAge Acceleration (**Figure 2C-D; Supplemental Table 9**, significant after FDR correction).

**Figure 2.**
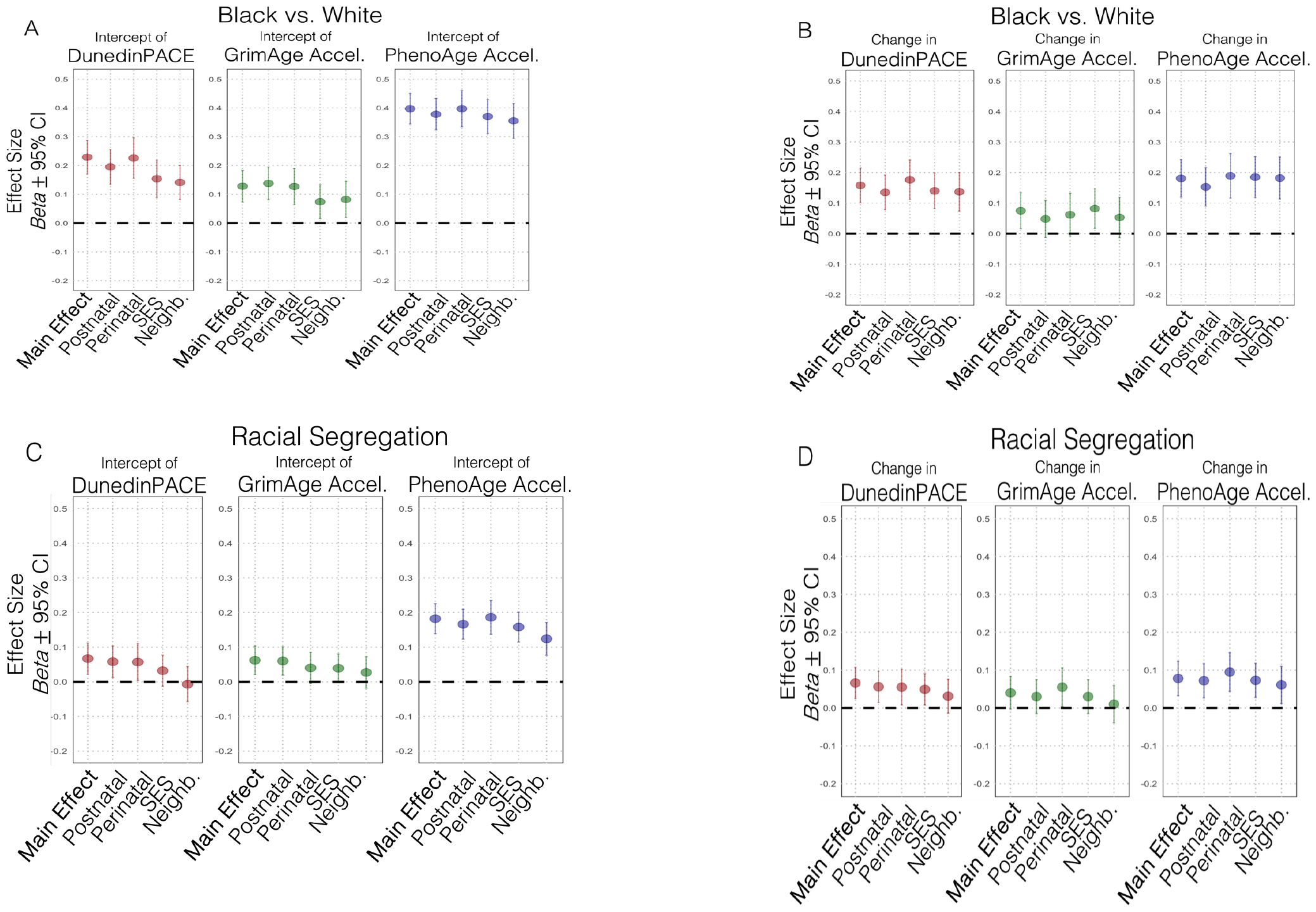
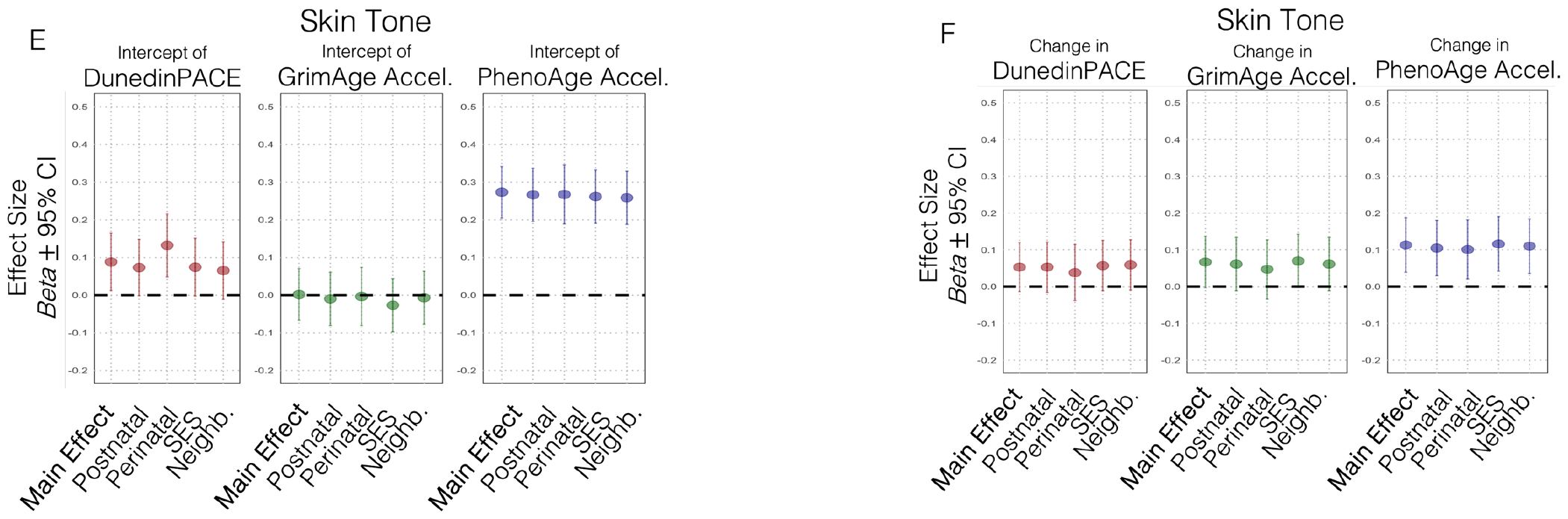
Racial and ethnic disparities in DNA-methylation measures of biological aging across adolescence. Panel. **A** depicts the difference in biological aging age-9 intercepts (DunedinPACE, GrimAge Acceleration, PhenoAge Acceleration) between Black compared to White identifying children without (i.e., Main Effect) and with covariate adjustment (covariates: postnatal factors, perinatal factors, family socioeconomic status [SES], and neighborhood disadvantage [Neighb.]). **Panel B** depicts the difference in longitudinal change from age-9-to-15 in biological aging between Black compared to White children without and with covariate adjustment. **Panel C** and **D** depict associations of neighborhood racial segregation with biological aging intercepts and longitudinal change, respectively. **Panel E** and **F** depict associations of skin tone with biological aging intercepts and longitudinal change, respectively.

**Figure 3.**
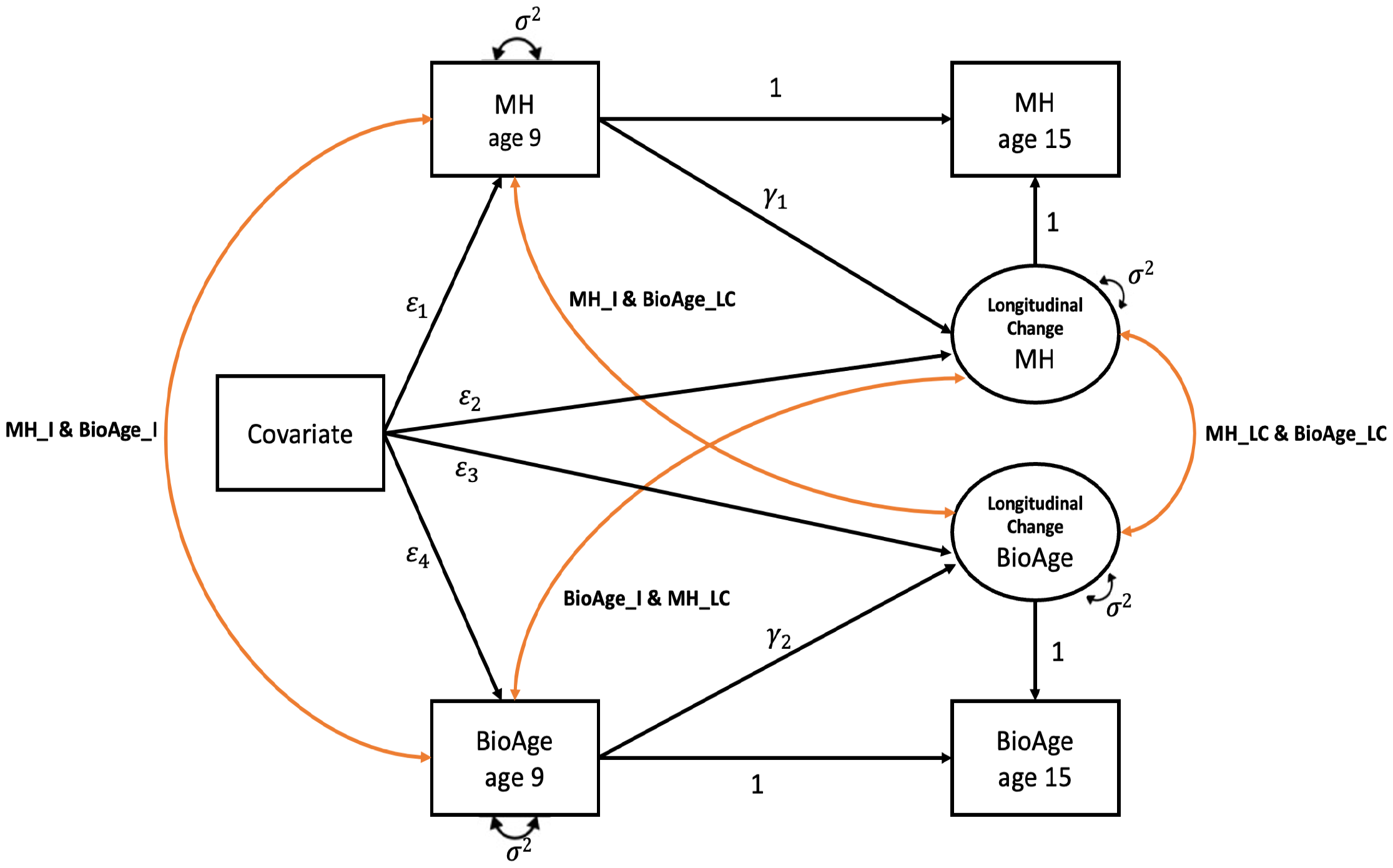
Bivariate latent change score model of mental health and DNA-methylation measures of biological aging across adolescence. Squares represent observed variables. Circles represent latent factors. Single headed arrows denote regressions and double headed arrows denote correlations. MH = Mental health measures of parent-reported externalizing and internalizing behaviors. BioAge = DNA-methylation measures of biological aging. I = Intercept. LC = Longitudinal change from 9-to-15-years. The estimated means of intercepts, longitudinal change, and covariates (*e*.*g*., prenatal factors) are not illustrated here, but were included in the model.

Amongst Black, Latinx and Multiracial children, darker skin tone was associated with a faster pace of aging intercept and more advanced biological age intercept and higher longitudinal increase in biological age as indicated by PhenoAge Acceleration (**Supplemental Table 10, Figure 2E-F**, significant after FDR correction). No significant associations were found for skin tone and GrimAge Acceleration (**Supplemental Table 10**).

Subsequently, we tested to what extent racial/ethnic disparities in biological aging were statistically accounted for by covariate adjustment for factors previously associated with DNAm measures of biological aging and/or structural racialization such as postnatal covariates (BMI, smoking, puberty status), perinatal birth factors (gestational age, birthweight, substance use during pregnancy), as well as proximal contextual factors (family socioeconomic status, neighborhood disadvantage, parental stress, parental closeness, and police interactions) The difference in biological aging between Black and White youth in biological aging largely persisted after accounting for postnatal covariates, perinatal birth factors, as well as proximal contextual factors (**Figure 2A & B, Supplemental Table 8**). Associations of neighborhood racial segregation with biological aging largely persisted after accounting for postnatal, perinatal, police and parenting factors, but associations with DunedinPACE and GrimAge Acceleration were fully accounted for by socioeconomic status and neighborhood disadvantage (**Figure 2C-D, Supplemental Table 9**). Similarly, associations of skin tone with biological aging largely persisted after accounting for postnatal, perinatal, police and parenting factors, and associations with PhenoAge Acceleration also remained significant after accounting for socioeconomic status and neighborhood disadvantage (**Supplemental Table 10, Figure 2E-F**).

### (3) Associations of mental health and biological aging

First, we examined whether longitudinal change from age-9-to-15-years in externalizing behaviors was associated with longitudinal changes from age-9-to-15-years in biological aging (see **Figure 3** for a graphical model and **Table 1** for parameter estimates). We found that higher longitudinal increases in externalizing behavior were correlated with higher longitudinal increases in pace of aging and biological age acceleration (DunedinPACE: *r*=.06, 95% CI=.03 to .13, *p* <.01; GrimAge Acceleration: *r*=.06, 95%CI= .02 to .11, *p*<.01; PhenoAge Acceleration: *r*=.05, 95%CI= .01 to .10, *p*<.05; significant after FDR correction). These associations largely persisted after accounting for perinatal and postnatal covariates as well as self-identified race/ethnicity and racial segregation (**Supplemental Table 11**; see **Supplemental Table 12** for longitudinal correlations from subgroup analysis of White, Black and Latinx groups).

Second, we tested whether longitudinal changes in internalizing behaviors were associated with longitudinal changes in biological aging. Higher longitudinal increases in internalizing behavior were correlated with higher longitudinal increases in pace of aging and biological age acceleration (DunedinPACE: *r*=.06, 95%CI=.01 to .10, *p* <.05, PhenoAge Acceleration: *r*=.06, 95%CI= .01 to .10, *p*<.05, significant after FDR correction; GrimAge Acceleration: *r*=.04, 95%CI= .01 to .08, *p*=.08). These associations largely persisted after accounting for postnatal covariates and perinatal birth factors as well as self-identified race/ethnicity and racial segregation (**Supplemental Table 11**).

Lastly, we tested if age-15 biological aging was associated with mental health from early childhood through adolescence. We found more advanced biological age at age 15 years, as indicated by GrimAge and PhenoAge Acceleration, was associated with a higher externalizing intercept (GrimAge: b= .12, 95%CI = .07 to .18, *p*<.01; PhenoAge: b= .07, CI = .13 to .12, p<.05), stronger decrease in childhood externalizing (GrimAge b= -.14, 95%CI = -.23 to -.04, *p*<.01), and a subsequently stronger increase over adolescence (GrimAge b= -.20, 95%CI = -.06 to -.03, *p*<.01), which remained significant after FDR correction. While the association with PhenoAge Acceleration was fully accounted for by socioeconomic variables, the association with GrimAge Acceleration largely remained significant after accounting for covariates (**Supplemental Table 13**). More advanced biological age, as indicated by GrimAge Acceleration, was also correlated with a higher internalizing intercept (b= .08, 95%CI = .02 to .15, *p*<.05; significant after FDR correction). This association was largely accounted for by postnatal covariates as well as family and neighborhood socioeconomic factors (**Supplemental Table 14**). A faster DunedinPACE-pace of aging at age-15-years was associated with higher concurrent levels of anxiety and depressive symptoms, and the association with anxiety symptoms remained significant after FDR correction as well as after covariate controls (anxiety: b= .07, 95%CI = .02 to .11, *p*<.01; depression: b= .05, 95%CI = .01 to .09, *p*<.05, **Supplemental Table 15**).

## Discussion

We leveraged a prospective birth cohort study to examine whether the emergence of racial disparities in mental health is linked to the emergence of racial disparities in DNAm measures of biological aging across childhood and adolescence. We find that children identifying as part of a racially marginalized group and children living in more racially segregated neighborhoods had higher levels of both externalizing and internalizing behaviors. Racial differences in externalizing behaviors were most pronounced for Black compared to White children, whereas all racially marginalized children had higher levels of internalizing symptoms than their White peers. Our findings align with previous studies on racial disparities in internalizing and externalizing behaviors in children, adolescents, and psychopathology in adults (Bernard et al., 2021; Del Toro et al., 2022; Mitchell et al., 2011, 2015; Wiggins et al., 2015). Moreover, amongst marginalized youth, we find that exposure to colorism, as indicated by darker skin tone, was associated with higher externalizing, but not internalizing, levels. Thus, our results substantiate neighborhood segregation and skin tone as mental health-relevant measures of racialization, including social hierarchies associated with colorism that privileges lighter skin shades over darker ones (Dixon & Telles, 2017; Hunter, 2007).

Additionally, Black compared to White identifying children, children living in more racially segregated neighborhoods, and marginalized children with darker skin tones, tended to have higher age-9 levels of biological aging and more biological age acceleration over adolescence. This is in line with previous cross-sectional findings in children and adults (Martz et al., 2024; Mitchell et al., 2016; Raffington, Schneper, et al., 2023b; Shen et al., 2023). For example, Hicken and colleagues (2023) find that Black compared to White identifying adults have higher GrimAge and PhenoAge Acceleration (GrimAge Acceleration: b = .42, 95% CI .20 to .64, *p*<.001; PhenoAge Acceleration: b = .29, 95% CI .02 to .57, *p*<.001). Notably, our saliva-based results of Black versus White disparities in 9-year-old children’s biological aging are partially of a similar magnitude to reports in adults (GrimAge Acceleration: b = .13, 95% CI .07 to .18, *p*<.001; PhenoAge Acceleration: b = .40, 95% CI .34 to .45 *p*<.001). Moreover, our longitudinal analyses show that racial disparities in biological aging increase over the course of adolescence, potentially highlighting a sensitive developmental period for lifespan health trajectories (Graf et al., 2022; Sawyer et al., 2012).

Even more so, these longitudinal increases in biological aging across adolescence were correlated with increases in internalizing and externalizing behavior. This is consistent with the interpretation that lower well-being has negative physical health consequences and vice versa (Kim et al., 2023; Prince et al., 2007). Alternatively, other factors, such as heightened daily life stress and vigilance stemming from ongoing racialization, could concurrently influence both within-person change in mental health and biological aging over adolescence (Castro-Ramirez et al., 2021; Goosby et al., 2018). Over time, an increased mental health burden could contribute to racial disparities in disease and mortality, alongside unequal access to healthcare and educational opportunities (Hicken et al., 2023). Importantly, effects of stress on DNAm measures of biological aging have been shown to be reversible if the stressor is removed (Poganik et al., 2023).

We further considered to what extent racial disparities were statistically accounted for by factors previously associated with structural racialization and/or DNAm, such as postnatal covariates, perinatal birth factors, as well as proximal contextual factors. Racial and ethnic disparities in mental health were largely statistically accounted for by socioeconomic variables, whereas disparities in biological aging were reduced, but remained visible, after statistically accounting for perinatal and postnatal covariates. Similarly, disparities of neighborhood segregation and colorism were largely statistically accounted for by socioeconomic factors. In contrast, measures of police interactions or parenting did not statistically account for racial disparities.

Structural racialization results in socioeconomic advantages for some racial groups and disadvantages for others (O’Brien et al., 2020). Accordingly, racially marginalized children were substantially more likely to reside in socioeconomically under resourced families and neighborhoods: Black and Latinx children were 82.8% and 43.2% more likely to reside in disadvantaged neighborhoods compared to White children, respectively (**see Supplemental Material Figure 1**). Hence, it is statistically challenging – and perhaps theoretically futile – to attempt to disentangle racialized and socioeconomic inequality in racially stratified population cohorts. Instead, advancements in our understanding of racialization and health will be made by applying intersectional perspectives and collecting dynamic measurements (*e*.*g*., economic health benefits differ across racial groups; measures of experienced racial discrimination are lacking; Collins et al., 2021). Regular exposure to discriminatory policies and actions, especially in low-income, racially segregated areas, contribute to the emergence of racial disparities in physiological and psychological burden (Clark et al., 2022; Paradies et al., 2015; Williams et al., 2019).

By applying DNA-methylation algorithms developed in adult studies of multi-system health and mortality to children, our study finds that the link between mental and physical health - both of which are racialized – emerges in the first two decades of life. The early onset of these racial differences underscores the need to assess behavioral and psychological manifestations, along with harmful social conditions, earlier in life alongside DNAm measurements. Future research with repeated DNAm measures from birth can explore how early in life these associations first become apparent and would benefit from more comprehensive and diverse measures of racialization. Programs promoting racial health equity must address the psychological and physical impacts of structural racialization in children and adolescents.

## Methods

### Participants

The Future of Families and Child Wellbeing Study (FFCWS) follows a sample of 4,898 children born in large US cities during 1998-2000. FFCWS oversampled children born to unmarried parents and interviewed parents at birth and ages 1, 3, 5, 9 and 15. During home visits, saliva DNA was collected the Illumina 450K and EPIC methylation arrays with ages 9 and 15 assayed on the same plate. DNAm data is available at age 9 (n=1971) and age 15 (n=1974). DNAm study participants self-identified race/ethnicity defined by study protocol as African-American/Black only (n=901, 47%), “Other” (n=52, 3%), Hispanic/Latinx (n=511, 26%), Multiracial (n=99, 5%), White (n=366, 19%). The University of Michigan and Princeton University Institutional Review board granted ethical approval. Informed written consent was obtained from study participants’ legal guardians in both cohorts. See **Table 2** for description of study measures and **Supplemental Table 1** for descriptives and for correlations between measures of interest.

## Measures

### Preprocessing DNA data

DNA extraction and methylation profiling for FFCW was conducted by the Notterman Lab of Princeton University and the Pennsylvania State University College of Medicine Genome Sciences Center. Due to the timing of assay completion 40% of the FFCW saliva samples were completed using the Illumina 450K chip and the remaining 60% used the Illumina EPIC chip. Methods for the two chips were standardized as much as possible, but all analyses were run separately for 450 and EPIC and then meta-analyzed. 450K DNAm image data were processed in R statistical software (4.1) using the Enmix package (Xu et al., 2016).

The red and green image pairs (n_samples_ =1811) were read into R and the Enmix preprocessENmix and rcp functions were used to normalize dye bias, apply background correction, and adjust for probe-type bias. The majority of sample filtering was applied using the ewastools packages (Heiss & Just, 2019). We dropped samples using the following criteria: if >10% of DNA-methylation sites had detection p-value >0.01 (n_samples_ =34), if there was sex discordance between DNAm predicted sex and recorded sex (n_samples_ =11), or if two sequential samples from the same individual exhibited genetic discordance between visits (n_samples_ =27). ENmix QCinfo function identified samples with outlier methylation values which were cut (n_samples_ =6). Technical replicates were removed n_samples_ =49). This gave us our final analytic sample (n=1684). DNAm sites were removed if they had detection p-value >0.01 in 5% of samples (n=33,376). Relative proportions of immune and epithelial cell types were estimated from DNAm measures using a childhood saliva reference panel (Middleton et al., 2022). EPIC DNAm image data were processed in R statistical software (4.1) using the ENmix package (Xu et al., 2016). The red and green image pairs (n_samples_ =2558) were read into R and the ENmix preprocessENmix and rcp functions were used to normalize dye bias, apply background correction, and adjust for probe-type bias. The majority of sample filtering was applied using the Ewastools packages ^7^. We dropped samples using the following criteria: if >10% of DNAm sites had detection p-value >0.05 (n_samples_=63), if there was sex discordance between DNA-methylation predicted sex and recorded sex (n=12), or if two sequential samples from the same individual exhibited genetic discordance between visits (n=30). ENmix QCinfo function identified samples with outlier methylation values which were cut (n=1) or samples that failed bisulfite conversion (n_samples_=7). Technical replicates were removed (n=168). This gave us our final analytic sample (n_samples_=2277). DNAm sites were removed if they had detection p-value >0.05 in 5% of samples (n=127,275). Relative proportions of immune and epithelial cell types were estimated from DNAm measures using a childhood saliva reference panel (Middleton et al., 2022).

## Supporting information

Supplemenatl Tabels

## Funding Acknowledgements

During her work on this paper, MA was a pre-doctoral fellow of the International Max Planck Research School on the Life Course (LIFE, www.imprs-life.mpg.de; participating institutions: Max Planck Institute for Human Development, Freie Universita Berlin, Humboldt-Universita zu Berlin, University of Michigan, University of Virginia, University of Zurich). This study was supported by the National Institute of Minority Health and Health Disparities under award numbers R01MD011716 and R01MD011716; the National Institute on Aging under award numbers R25AG05322 ; the National Institute of Child Health and Human Development under award R01HD036916, R01HD076592, and P2CHD042849; and the National Institute of Mental Health under award number R01MH103761. LR is faculty member at the International Max Planck Research School on the Life Course and received funding from the Max Planck Society. The funding bodies had no role in the design, collection, analysis or interpretation of the study.

