## Supplemental Material for "Linked emergence of racial disparities in mental health and epigenetic biological aging across childhood and adolescence"

### Supplemental Material Table 1

Table 1. List of preregistered analyses (see https://osf.io/5sejf/), deviations, and results if not reported in main text.

| Research Question | Preregistered analysis | Deviation from preregistration, if applicable | Reporting of the result / result if not reported in the main text |
|---|---|---|---|
| *1)* Associations of race and mental health | | | |
| Are children's race/ethnicity and neighborhood racial segregation associated with mental health? Is this association moderated by socioeconomic status? | We use latent growth curve models of internalizing/externalizing intercepts (age 3 year) and slopes (change over time from 3 to 15 years) as well as in cross-sectional analyses of age-15 anxiety and depression. | | See result section and Supplemental Table 3, 4 (for longitudinal analyses) and Supplemental Table 7 (for cross-sectional analyses). |
| | Regress mental health on children's race/ethnicity. | | See result section and Supplemental Table 3 and 4 (for longitudinal analyses) and Supplemental Table 7 (for cross-sectional analyses). |
| | Regress mental health on children's race/ethnicity, socioeconomic contexts (family level SES and neighborhood disadvantage), and an interaction of children's race/ethnicity by socioeconomic contexts. | In the pre-registration, we refer to socioeconomic contexts for both family and neighborhood SES. In our manuscript, we call this family level SES and neighborhood disadvantage to enhance clarity. | See result section and Supplemental Table 3 and 4. There were no significant neighborhood *race/ethnicity interactions, there was a significant family level SES* race interaction, for externalizing of LatinX Children. See Supplemental Table 3. |
| | Regress mental health on neighborhood racial segregation. | | See result section and Supplemental Table 5. |
| | Regress mental health on neighborhood racial segregation, socioeconomic contexts (family level SES and neighborhood disadvantage), and an interaction of neighborhood racial segregation by socioeconomic contexts (family level SES and neighborhood disadvantage). | In the pre-registration we refer to racial context, in the manuscript to neighborhood racial segregation to enhance clarity. | There were no significant interaction effects of racial contexts by family or neighborhood SES for internalizing and externalizing. |
| To what extent do police interactions and socioeconomic contexts account for racial/ethnic disparities in mental health? | For significant associations, regress mental health on children's race/ethnicity and police interactions. | | See result section and Supplemental Table 3 and 4. |

| | | | |
|---|---|---|---|
| | For significant associations, regress mental health on children's race/ethnicity and socioeconomic contexts (family level SES and neighborhood disadvantage), | | See result section and **Supplemental Table 3 and 4.** |
| Is skin tone associated with mental health amongst racialized youths? Is this association moderated by race/ethnicity? | For African-American/Black, Hispanic/Latinx, and Multiracial participants only, regress mental health on skin tone. | | See result section and **Supplemental Table 6.** |
| | For African-American/Black, Hispanic/Latinx, and Multiracial participants only, regress mental health on skin tone, race/ethnicity (African-American/Black, Hispanic/Latinx, and Multiracial) and an interaction of skin tone by race/ ethnicity. | | There were no significant interaction effects of skin tone by race/ethnicity on mental health. |
| Does parenting moderate associations of racial/ethnicity with mental health? | Regress mental health on children's race/ethnicity and parenting and an interaction of parenting by race/ethnicity. | | See result section and **Supplemental Table 3 and 4**. Only significant interaction for externalizing; race by closeness for Multiracial Slope 2 (b = -1,024, CI =-1,861 to -0,187, p=0,016) |
| | Regress mental health on neighborhood racial segregation, parenting, and an interaction of race by parenting. | | See result section and **Supplemental Table 5**. There were no significant interaction effects of neighborhood racial segregation by parenting on mental health. |
| Does gender moderate associations of race/ethnicity with mental health? | Regress mental health on children's race/ethnicity, gender, and an interaction of race by gender. | | There was a significant race*sex interaction for Black children on Slope 1 and Slope 2 for Externalizing and Internalizing behavior, with more prominent effects for Black Boys (See results section and **Supplemental Table 4).** |
| | Regress mental health on neighborhood racial segregation, gender, and an interaction of neighborhood racial segregation by gender. | | There were no significant racial segregation * sex interactions. |
| *2)* Associations of race and DNAm-aging | | | |
| Are race/ethnicity associated with DNAm-aging measured at age | Regress DNAm-aging on children's race/ethnicity, separately for ages 9 and 15. | Instead of modeling cross-sectional effects, we applied a latent | See result section and **Supplemental Table 8.** |

| | | | |
|---|---|---|---|
| 9 and age 15? Is this association moderated by socioeconomic contexts? | | change score to investigate the effects of race/ethnicity on both the intercept and the change (i.e., delta). We deviated from the preregistration as this analysis allowed us to include the strength of the longitudinal data, and to gain better insights into developmental patterns. | |
| | Regress DNAm-aging on children's race/ethnicity, socioeconomic contexts (family level SES and neighborhood disadvantage), and an interaction of children's race/ethnicity by socioeconomic contexts, separately for ages 9 and 15. | See above | See result section and Supplemental Table 8. There was no significant Family SES* race interaction on biological aging measures. There was a significant neighborhood disadvantage*race interaction on the intercept of GrimAge acceleration and the longitudinal change of DunedinPACE (see **Supplemental Table 8,** only the interaction with the longitudinal change of Dunedinpace was significant after FDR correction). The association between neighborhood disadvantage and biological aging was more prominent for White children than for marginalized children, although this did not hold FDR correction (intercept GrimAge Acceleration: White b=.19, 95%CI=.10 to .27, p<.001, Black b=.04, 95%CI=-.03, .17, p=.25, Latinx b=.07, 95%CI=-.01, .15, p=.08; delta DunedinPACE from age 9 to 15: White b=.27, 95%CI=.18 to .36, p<.001, Black b=.02, 95%CI=-.04, .08, p=.51, LatinX b=-.01, 95%CI=-.09, .07, p=.78). |

| | | | |
|---|---|---|---|
| | | | This may be driven by the fact that racially marginalized children are far more likely to live in socioeconomically under resourced neighborhoods (see **Supplemental Material Figure 1**). |
| | Regress DNAm-aging on neighborhood racial segregation, separately for ages 9 and 15. | See above | See result section and **Supplemental Table 9**. |
| | Regress DNAm-aging on neighborhood racial segregation, socioeconomic contexts (family level SES and neighborhood disadvantage), and their interaction of neighborhood racial segregation by socioeconomic contexts, separately for ages 9 and 15. | See above | There were no significant interaction effects of SES*neighborhood racial segregation on biological aging measures. |
| To what extent do socioeconomic contexts and police interactions account for racial/ethnic disparities in DNAm-aging? | For significant associations, regress DNAm-aging on children's race/ethnicity and police interactions. | See above | See **Supplemental Table 8**. We did not see significant association between police interactions and biological aging measures. |
| | For significant associations, regress DNAm-aging on children's race/ethnicity and socioeconomic contexts (family level SES and neighborhood disadvantage), | | See **Supplemental Table 8**. |
| Is skin tone associated with DNAm-aging within racialized youths? | For African-American/Black, Hispanic/Latinx, and Multiracial participants only, regress DNAm-aging at age 9 and 15 on skin tone, race/ethnicity (African-American/Black, Hispanic/Latinx, and Multiracial) and a skin tone x race interaction. | | See result section, and **Supplemental Table 10**. There were no significant skin tone*ethnicity/race interactions on biological aging measures. |
| Does parenting moderate associations of socioeconomic contexts and police interactions on DNAm -aging? | For significant associations, regress DNAm-aging on socioeconomic contexts and parenting. | See above | Associations between parenting and DNAm aging were not significant, see **Supplemental Table 8, 9** and **10**. There were no significant interaction effects for parenting*SES. |
| | For significant associations, regress DNAm-aging on police interaction and parenting. | | There were no significant interaction effects for parenting*police on biological aging. |
| Does gender moderate associations of | Regress DNAm on children's race/ethinicty, gender, and an interaction of race by gender. | | While there was a main effect of sex on DNAm measures (see |

| | | | |
|---|---|---|---|
| race/ethnicity with mental health? | | | Supplemental Table 8, 9, 10), there was no significant interaction effect of race by sex on biological aging. |
| Is this association accounted for by covariates? | For significant associations, we rerun analyses adding covariates. | While we did not pre-register to include perinatal covariates, we decided to include them as they are associated with measures of biological aging. | For all significant associations, we ran covariate analyses including postnatal (BMI, smoking, puberty status), and perinatal birth factors (gestational age, birthweight, substance use during pregnancy). See Supplemental Table 8-10. |
| Are mental health trajectories across childhood associated with DNAm-aging in adolescence? | Regress DNAm-aging at age 15 on internalizing intercept and slope. | | See result section and Supplemental Table 14. |
| | Regress DNAm-aging at age 15 on externalizing intercept and slope. | | See result section and Supplemental Table 13. |
| | Regress DNAm-aging at age 15 on anxiety | | See result section and Supplemental Table 15. |
| | Regress DNAm-aging at age 15 on depression | | See result section and Supplemental Table 15. |
| Does DNAm-aging at age 9 predict externalizing and internalizing at age 15 and vice versa? | We will fit a fixed-effects regression model to examine the correlation of within-person changes in externalizing and internalizing with changes in DNAm-aging from age 9 to 15 years. | Instead of applying a fixed-effects regression model (FE) and a cross-lagged panel model (CLPM), we applied a latent change model which combines the features of CLPM and FE in one model allowing to more precisely investigate the effect of changes in one variable on changes in the other variable. (McArdle, 2009; Mund & Nestler, 2019) | See results section Table 1, and Supplemental Table 11 |
| | We will fit a bivariate random intercept cross-lagged panel model to test whether internalizing and externalizing behavior at age 9 predict DNAm-aging at age 15 and vice versa. | See answer above. | See answer above. |
| Is this association accounted for by covariates? | For significant associations, we rerun analyses adding covariates. | While we did not pre-register to include perinatal covariates, we decided to include them as they are associated | For all significant associations, we ran covariate analyses including postnatal (BMI, smoking, puberty status), and perinatal birth factors |

| | | | |
|---|---|---|---|
| | | with measures of biological aging. | (gestational age, birthweight, substance use during pregnancy) as covariates. See results section and **Supplemental Table 11 and 12**. |
| All models will include age, gender and an age -by-gender interaction term. | | We utilized data from a cohort study assessing children at the same age. We therefore only included gender, but not age or age by gender as interaction term in our analyses. | |

### Supplemental Material Figure

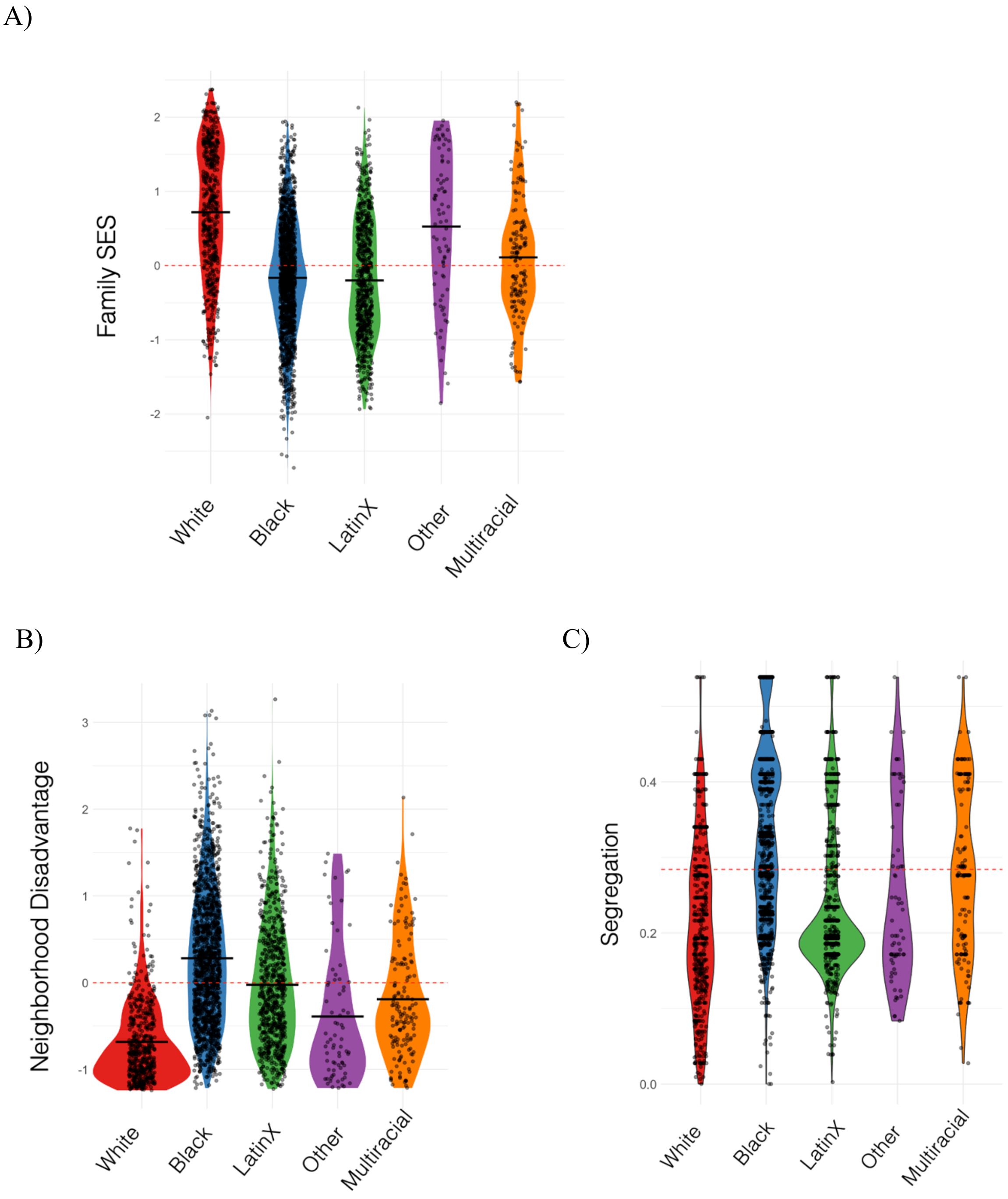

**Supplemental Figure 1. Racial and ethnic disparities in socioeconomic status, neighborhood disadvantage and neighborhood segregation. Panel A** plots family socioeconomic status (SES) by racial/ethnic identities, where higher scores indicate higher family SES. **Panel B** plots neighborhood disadvantage by racial/ethnic identities, where higher scores indicate more neighborhood disadvantage. **Panel C** plots neighborhood segregation by racial/ethnic identities, where higher scores indicate more segregation.
